# Modelling low-dimensional interacting brain networks reveals organising principle in human cognition

**DOI:** 10.1101/2023.11.20.567824

**Authors:** Yonatan Sanz Perl, Sebastian Geli, Eider Pérez-Ordoyo, Lou Zonca, Sebastian Idesis, Jakub Vohryzek, Viktor K. Jirsa, Morten L. Kringelbach, Enzo Tagliazucchi, Gustavo Deco

## Abstract

The discovery of resting state networks shifted the focus from the role of local regions in cognitive tasks to the ongoing spontaneous dynamics in global networks. Recently, efforts have been invested to reduce the complexity of brain activity recordings through the application of nonlinear dimensionality reduction algorithms. Here, we investigate how the interaction between these networks emerges as an organising principle in human cognition. We combine deep variational auto-encoders with computational modelling to construct a dynamical model of brain networks fitted to the whole-brain dynamics measured with functional magnetic resonance imaging (fMRI). Crucially, this allows us to infer the interaction between these networks in resting state and 7 different cognitive tasks by determining the effective functional connectivity between networks. We found a high flexible reconfiguration of task-driven network interaction patterns and we demonstrate that can be used to classify different cognitive tasks. Importantly, compared to using all the nodes in a parcellation, we obtain better results by modelling the dynamics of interacting networks in both model and classification performance. These findings show the key causal role of manifolds as a fundamental organising principle of brain function, providing evidence that interacting networks are the computational engines brain during cognitive tasks.

## Introduction

The high dimensionality of brain activity recordings is one of the major obstacles hindering experimental and theoretical efforts in neuroscience. At the level of single cells, the human brain is constituted of billions of neurons and synapses where these neurons interact (Sporns et al., 2005); however, coarser scales can also be understood as complex interconnected systems where nodal dynamics represent the aggregate of a macroscopic population of neurons (Deco et al., 2008, 2011). Even if exhaustive recordings to these systems could be obtained, it would remain difficult to reach encompassing theoretical principles due to the sheer complexity of the data. In particular, addressing the behaviour of individual cells does not suffice for this endeavour, since the high interconnectivity of the brain facilitates the emergence of coordinated activity (Biswal et al., 1995; Deco et al., 2008), and thus of distributed representations where information is encoded in the global dynamics of large populations of neurons.

The discovery of resting state networks was a first step towards lowering the dimensionality of brain signals (Beckmann et al., 2005; Damoiseaux et al., 2006), which radically shifted the focus from the role of local regions in cognitive tasks to the ongoing spontaneous dynamics in global networks. In recent years, considerable efforts have been invested to tackle distributed representations by means of nonlinear methods capable of finding low dimensional representations of brain activity at multiple spatiotemporal scales, from cortical microcircuits (Chaudhuri et al., 2019; Mitchell-Heggs et al., 2022) to whole-brain dynamics measured with functional magnetic resonance imaging (fMRI) (Gao et al., 2021; Glomb et al., 2021; Luppi et al., 2023; Yonatan Sanz Perl et al., 2023; Rué-Queralt et al., 2021; Vidaurre et al., 2018; Vohryzek et al., 2023), among other neuroimaging methods. The success of this approach does not only rely on methodological considerations, but also on the fundamental characteristics of brain activity. Despite the very large number of degrees of freedom of brain dynamics, coordinated cognition and behaviour cannot exist without the integration of this activity (Shine et al., 2016; Tononi et al., 1998, 2016). Neural information processing also exhibits redundancy (Hennig et al., 2018; Shine et al., 2019; Tononi et al., 1999), where the activity of several cells is constrained to develop within an abstract geometrical space of lower dimensionality, generally known as *manifold*. The organization of brain activity into manifolds with a reduced number of dimensions, named networks or modes, is a frequently replicated phenomenon, and has been proposed as a fundamental aspect of brain dynamics, fulfilling different scale-dependent computational roles (Pang et al., 2016). The manifold spans a subspace of modes, in which the brain activity evolves in time following rules that determine the behaviour of the system on that manifold defining a flow. (Pillai & Jirsa, 2017). In other words, brain activity can be represented as a dynamical system in a low-dimensional space capturing the time evolution of the system in that space collapsing high dimensional information to the manifold.

The so-called structured flows on manifolds (SFMs) emerge from basic multi-scale processes such as symmetry breaking and near-criticality (Jirsa & Sheheitli, 2022). Indeed, a direct link has been proposed between the behavior and the flow of the low-dimensional manifold underlying the brain dynamics(Fousek et al., 2023; Jirsa, 2020). In particular, it has been shown that manifold coordinates are relevant describing cognitive brain processing and behaviour by parametrizing motor control (Gallego et al., 2017), perception (Chandak & Raman, 2023; Stringer et al., 2019), cognition and attention (Song et al., 2023), navigation (Derdikman & Moser, 2010) and sleep (Chaudhuri et al., 2019), among others examples. Overall, several investigations had mapped macroscales brain activity to low dimensional manifold representations linking measures of neural activity to cognition, however it is not clear how these networks interact and how these interactions are related with the cognitive processing.

Therefore, we directly investigated how the interaction between these low-dimensional networks emerge as an organising principle supporting cognitive function and we asked whether the reconfiguration of these interactions adequately tracks brain activity as the participants perform different tasks. To address this question, we first assume that brain activity is most adequately described in terms of a reduced number of abstract variables, here termed *networks*, each encoding the simultaneous dynamical behaviour of multiple macroscopic brain regions. These modes are neurobiologically meaningful as they emerge due to the large-scale anatomical and functional organization of the brain. Formally, modes are the coordinates spanning the subspace, in which the low-dimensional manifold is embedded. The *intrinsic dimensionality* of brain dynamics corresponds to the minimum number of such modes required for its description. Based on a previous works, we used a deep learning architecture known as variational autoencoders (VAE) to obtain low-dimensional representations of whole-brain activity from a large-scale fMRI data set of more than 1000 participants (Human Connectome Project, HCP). We modelled the dynamics of each coordinate on the manifold by a non-linear oscillator, representing the mode and operating close to the bifurcation point. The dynamical behaviour changes from fixed point dynamics towards self-sustained oscillations, as successfully applied in several prior works modelling the dynamics of the high dimensional state space (Deco et al., 2017; Jobst et al., 2017; Yonatan S Perl et al., 2022). To optimize the models, we inferred the effective connectivity between the modes considering the level of non-equilibrium dynamics of brain activity quantified by the non-reversibility of the signals (termed GCAT, following the work of Kringelbach et al. 2023 (Kringelbach et al., 2023)). We explored different manifold dimensions, and we demonstrated that by modelling brain dynamics on the manifold we achieve superior models compared to the traditional ones constructed in the source state space. Crucially, by investigating the reconfiguration of the interaction between networks across different tasks we were able to track brain activity more precisely during seven cognitive tasks included in the HCP dataset. These results establish that the inferred connectivity in the manifold space provided more informative results compared to those obtained solely in the source space of empirical recordings.

## Results

### Methodological overview

Our methodological approach encompasses two complementary stages addressing the main questions we posed. The first analysis involves assessing the optimal dimension of the reduced space which yield the bests computational dynamical models of interacting networks, and the second is focused on build these models to investigate how this interaction is modified during cognitive tasks. A summary of the proposed framework is displayed in **Figure 1**.

**Figure 1.**
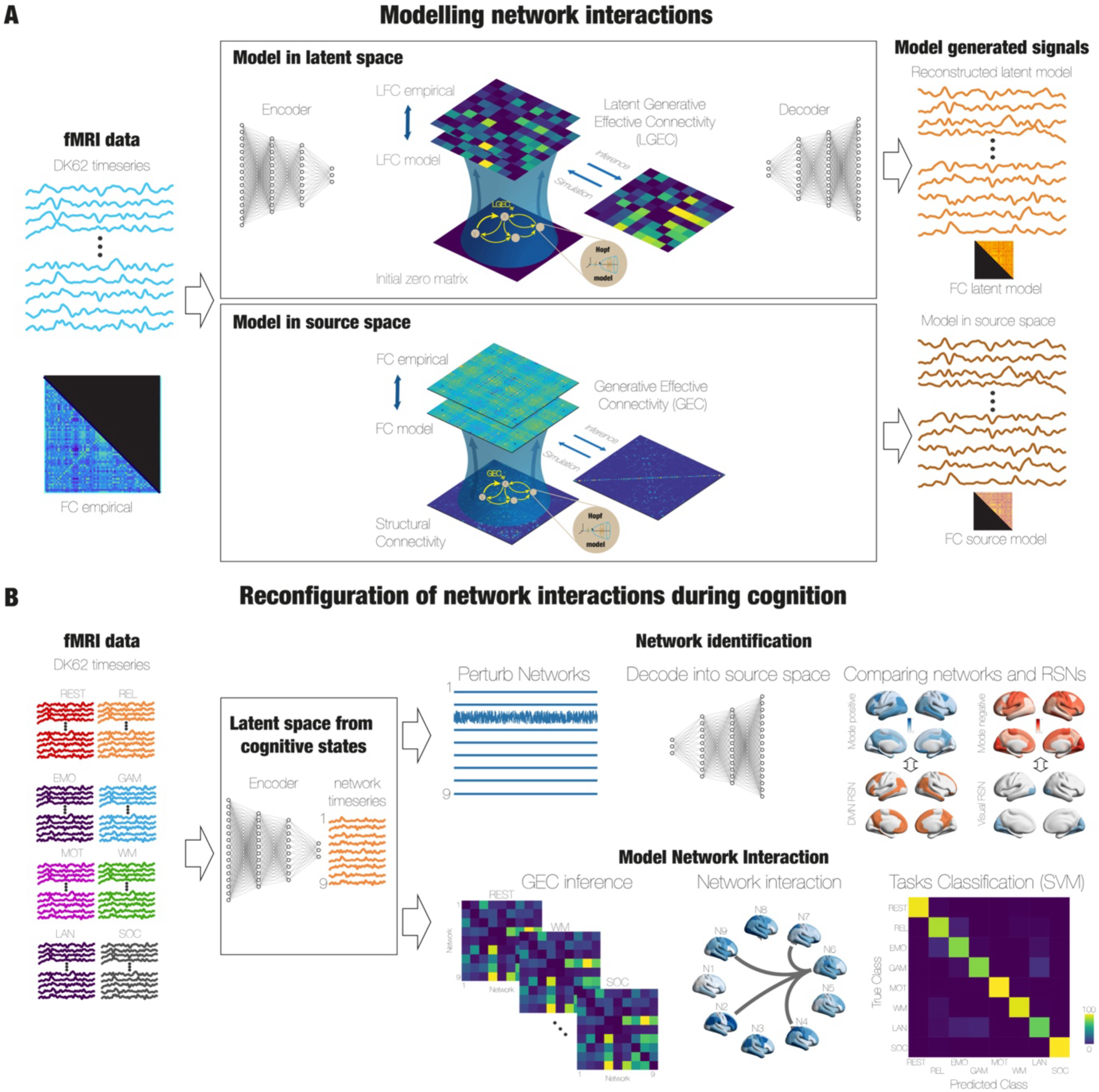
Overview of pipeline. **A)** We trained a variational AE (VAE) using the parcellation with N=62 brain regions, which consists of the same three structures, encoder, latent space and decoder, but also including a regularization loss function that gives VAEs generative properties. To model the dynamics of each network in the latent space, we utilized a non-linear Stuart-Landau oscillator near the bifurcation point, i.e. near the critical regime. In order to determine the connections between the latent variables, we employed the GCAT framework developed by Kringelbach and colleagues. We explored the performance of that models at different number of latent dimensions from 5 to 12. The result of this procedure is an optimized effective connectivity between the latent variables, LGEC, which captures the departure from detailed balance and generates the non-equilibrium dynamics observed in brain signals. We also created whole-brain models in the source space to fit the same empirical data by repeating the same optimisation procedure for the latent space mode, i.e., the inference of the GEC by implementing the GCAT framework. We compared the performance of both models and we found that models in the latent space more faithfully reproduce the empirical data quantified by the similarity between the empirical and modelled FC. **B)** We trained the (VAE) with nine dimensions in the latent space using the spatial patterns over time obtained from the combined fMRI data from 7 cognitive tasks from the HCP data set. We investigated which brain regions contributed to each of the nine latent modes previously found in terms of brain functional networks. To do so, we created a set of surrogate signals in the latent space by introducing standard Gaussian noise into one latent dimension while keeping the other eight dimensions devoid of any signal. We repeated by changing the noise signal for each of the nine modes. We then decoded the surrogate latent signals in each case obtaining the spatial pattern for each mode in source space. We then associate these patterns with the activation or deactivation of the seven resting state networks from Yeo. Finally, we assessed whether the interaction of these networks is driven by tasks activity by building computational models obtaining a network interaction matrix for each task. We found that these interactions are flexible, showing high variability across tasks, allowing us to train a high-performance classifier based on that information, surpassing the classification capability of models trained in the source space.

In the first stage (**Figure 1A**), we created dynamical models of the low-dimensional manifold obtained from the fMRI data of resting-state participants in the Human Connectome Project (HCP) in a cortical brain the DK62 cortical brain parcellation (Deco et al., 2021), which was constructed using the Mindboggle-modified Desikan–Killiany parcellation (Desikan et al., 2006), with a total of 62 cortical region. To do so, we used as input to the VAE the time-by-time spatial patterns from each region empirical timeseries (**Figure 1A**), i.e., the values of the *N* brain regions at each time, and we created a training set to optimize the parameters of the VAE. Importantly, a VAE consists of three key components: the encoder network, the middle layer (referred to as the bottleneck or latent space), and the decoder network. In particular, a VAE presents generative properties by forcing a regularization of the latent space through incorporation of a regularization term during the training process, the VAE ensures that the decoding step produces outputs that are both relevant and meaningful (Kingma & Welling, 2013) (see Methods). Notice that since our aim was to construct generative models in the latent space with the intention of investigating the decoded modelled signals, regularization of the latent space provided by VAEs was essential.

Subsequently, we constructed computational models in the latent space by describing the dynamics of each latent network using a non-linear Stuart-Landau oscillator close to the bifurcation point, i.e. close to the critical regime (**Figure 1A**). To infer the connectivity between the latent variables, we implemented the GCAT framework developed by Kringelbach and colleagues, which involves a gradient descent procedure where the effective connectivity is updated at each step, considering the disparities between the empirical and simulated functional connectivity, as well as the forward shifted connectivity, to generate a good fit of both observables (i.e. zero and one-lag functional connectivity) at the same time. The outcome of this procedure is an optimized effective connectivity between the latent variables (LGEC), which captures the breakdown of detailed balance, reproducing the non-equilibrium dynamical behaviour of brain signals(Kringelbach et al., 2023). We then explored the fitting performance of the computational models as a function of the latent dimension. Finally, we were able to decode the modelled signals to further generate observables that we compared with those computed in the original state space to quantify the performance of modelling the networks that constitute the low-dimensional manifold. We also created whole-brain models fitted to the same the empirical data in the source space, i.e., we replicated the GCAT framework in the original data consisting in the timeseries of each brain region (**Figure 1A**). We then compared the performance of both model in terms of the similarity between the empirical and modelled functional connectivity (FC). We found that the optimal dimension of latent model which more faithfully reproduces the empirical data is nine and that these models are better compared with the source space model.

In the second stage (**Figure 1B**), we trained the (VAE) using the spatial patterns over time obtained from the combined fMRI data from 7 cognitive tasks from the HCP data set. Based on the previous analysis we determined the dimensionality of the latent space in nine networks. To assess the relation within these networks and brain regions, we created a set of surrogate signals in the reduced space by introducing standard Gaussian noise into one network while keeping the other eight dimensions devoid of any signal. We repeated by changing the noise signal for each of the nine modes. We then decoded the surrogate signals in each case to obtain the corresponding spatial pattern in source space. We then associate these patterns with the activation (mode positive) or deactivation (mode negative) of the seven resting state networks from Yeo(Thomas Yeo et al., 2011) (**Figure 1B upper row**). Finally, to investigate how the interaction of these networks is tasks-driven activity reconfigured we built computational models obtaining a network interaction matrix for each task. We found that these interactions are flexible, showing high variability across tasks, allowing us to train a high-performance classifier based on that information, surpassing the classification capability of models trained in the source space (**Figure 1B lower row**).

### Models of interacting manifold networks outperform high dimensional models in the original space

We employed a VAE to obtain a low-dimensional manifold representation of fMRI data obtained from resting-state healthy participants from the HCP dataset. First, we determined the dimensionality of the low-dimensional manifold, i.e., number of networks, by assessing the reconstruction error of the autoencoder using three different grained brain parcellations. We found that, independent of the input dimensionality, the reconstruction error shows a significant decrease in the initial dimensions until it reaches an elbow point at approximately a latent dimension of 10, beyond which the error remains relatively constant (see **Supplementary Figure 1**). We also show in **Supplementary Figure 1** the derivative of the reconstruction error with respect to the latent dimension, demonstrating that for all three cases, around and above a dimension of 10, the value is close to zero. This confirms that no major changes in the reconstruction error occur at higher dimensions. It is worth mentioning that different parcellations show different absolute values of reconstruction error, which, like many other quantities, depend on the parcellation graining (e.g., graph theoretical measures (Domhof et al., 2021), structure function relationship (Messé, 2020), entropy (Kobeleva et al., 2021), in a recent work the reconstruction of an autoencoder (Jamison et al., 2024) and a systematic evaluation in fMRI (Luppi et al., 2024)) However, here we focused on the number of dimensions that stabilize the reconstruction error, independent of the absolute value of the error. We then explored in the following sections the optimal low-dimensional space dimension around 10 using a coarse-grained parcellation scheme comprising 62 regions (see Methods and Supplementary Material).

Subsequently, we modelled the dynamics of each network (i.e. mode) using non-linear Stuart-Landau oscillators, which exhibit dynamic characteristics determined by a Hopf bifurcation (see Methods for the mathematical expression of the model).

Typically, these models are fitted to reproduce empirical observables, such as the functional connectivity (FC) (Ipiña et al., 2020; Yonatan Sanz Perl et al., 2021). In our case we built a manifold model to fit the empirical functional connectivity between latent space modes (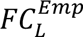 the subindex *L* stands for FC computed in the latent space and super index *emp* means that uses encoded empirical data), which is computed as the pairwise Pearson’s correlation between the latent modes, and the empirical latent forward-shifted connectivity ( 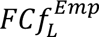), which represents the pairwise Pearson’s correlation between the signals in latent space and the same signals shifted forward in time (see Methods). To improve the model fitting capacities, we averaged the 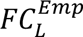 and 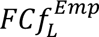 by subgroups of 10 participants and created 100 models that fit these average matrices.

To construct the model of the low-dimensional manifold, we fixed the bifurcation parameter for each mode to *a_j_* = −0.02 and we built the whole manifold model by coupling the oscillators representing each latent neural mode. The connectivity between the modes was inferred using an iteratively pseudo-gradient algorithm, which was proposed by Kringelbach and colleagues for models in the source space (Kringelbach et al., 2023). Briefly, this procedure consists in an iterative optimization where in each step the connections are updated considering the differences between the empirical and the modeled functional connectivity in the latent space (shifted and non-shifted) weighted by a learning factor ς (see Methods). This iterative process enabled us to accurately capture the underlying relationships between latent modes. By fitting the 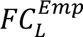 at the same time that 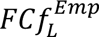, we could effectively capture the non-equilibrium dynamic patterns in latent signals (Kringelbach et al., 2023). The result of this process is an enhanced effective connectivity for each one of the 100 models between the latent modes (LGEC), which also captures the disruption of the detailed balance, leading to the generation of non-equilibrium dynamic patterns in latent signals. We show that this disrupted detailed balance is also captured by the decoded signals.

We proceeded to assess the overall performance of the complete framework by examining the similarity between the modelled and decoded FC (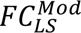 computed as the Pearson’s correlation of the signals modelled in the low-dimensional manifold and then decoded to the source space) and the empirical FC in the source high dimensional space (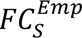) quantified by the correlation between both FC. We also computed the correlation between the forward matrices 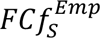 and 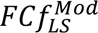 (**Figure 2A**). Both metrics exhibited similar behaviour, displaying an optimal point at latent dimension 9, with decreasing performance observed for higher and lower dimensions. To validate the reliability of our approach, we constructed the model directly in the source state space using the GCAT framework, following the exact same methodology as employed in the latent space. However, we observed significantly lower levels of fitting for both observables compared to the results obtained from the low-dimensional manifold modelling (blue boxplot in Figure 2A). This discrepancy was observed not only for the optimal dimension but also across all dimensions under evaluation, demonstrating that even outside of the optimal manifold representation, the low-dimension representation is still more accurate than the high-dimensional original space. Importantly, by modeling the low-dimensional space, we are able to improve the quality of the fitting, avoiding the need to force the model to reproduce redundant and noisy information present in the high-dimensional representation. Consequently, our low-dimensional models accurately capture the interaction between these networks as a signature of cognitive states, allowing for increased performance of the SVM classifier as we demonstrate later.

**Figure 2.**
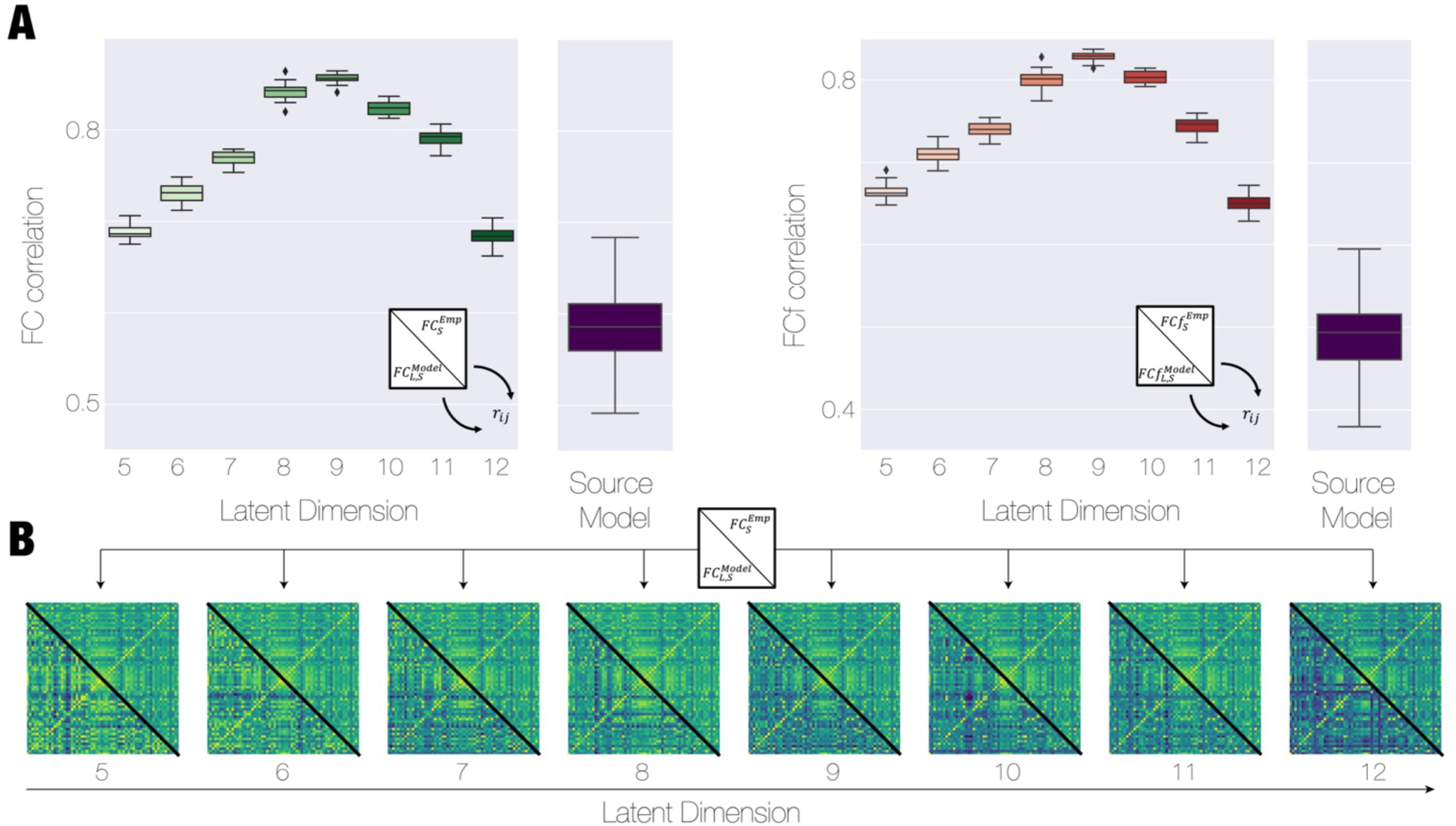
Modelling the low-dimensional brain manifold reconstructs empirical data, outperforming models in the original state space. **A**) We generated with a VAE a low-dimensional manifold representation of fMRI data from resting-state healthy participants in the Human Connectome Project (HCP) using a coarse-grained parcellation scheme consisting of 62 regions. To model the dynamics of each latent dimension within the manifold, we employed non-linear Stuart-Landau oscillators. These oscillators exhibit dynamic characteristics that are governed by a Hopf bifurcation. We constructed a manifold modes model that accurately fit the 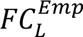 and 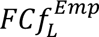 which represent the pairwise Pearson’s correlation between the signals in the latent space and the same signals with the corresponding shifted forward in time, respectively. The results of the performance of the complete framework in terms of reconstruction of 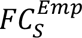 and 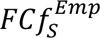 are quantified as the correlation between these matrices and the ones obtained through decoding the manifold models for each dimension. The maximum correlations are reach for latent dimension 9 and decreasing both for low and high dimension. Importantly, we demonstrated that these models overcome the performance of a model developed in the high-dimensional source space (blue boxplot), not only for the optimal dimensions but also for all latent dimension explored**. B)** The hybrid matrices display in the upper diagonal triangle the 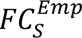 and in the lower diagonal triangle the 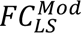 to observe the similarity between both matrices when explore different latent dimension ranging from 5 to 12.

Figure 2B depicts hybrid matrices as a function of the latent dimension that compare the empirical functional connectivity 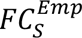 (upper triangle) and one obtained as 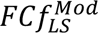 (lower triangle).

### Source network identification of tasks-activity latent networks

We investigated how cognitive processing is related to dynamics in the low-dimensional networks by analysing fMRI data from 7 cognitive tasks from the HCP data set (Social [SOC], Language [LAN], Working Memory [WM], Motor [MOT], Gambling [GAM], Emotion [EMO], Relational [REL]). Similar to the previous section, we trained the Variational Autoencoder (VAE) using the spatial patterns over time obtained from the combined data of the rest condition and the seven tasks (see Methods). We fixed the latent dimension in nine based on the previous results and we investigated which brain regions contributed to each of the nine latent networks in terms of brain functional networks. To assess this, we obtained one spatial pattern on the empirical original space associated to each latent mode (see Methods). We evaluated these patterns in terms of the reference functional brain networks estimated by Yeo and colleagues: Visual (VIS), Somatomotor (SM), Dorsal Attention (DA), Ventral Attention (VA), Limbic (Lim), Frontoparietal (FP) and Default mode (DMN), known as the 7 resting-state networks (RSNs) from Yeo(Thomas Yeo et al., 2011). We computed the percentage of participation of each brain region within the DK62 parcellation to each of the 7 RSNs. We then computed the Pearson’s correlation between each of the nine patterns corresponding to each neural mode and the percentage of participation of each brain region to each RSN. In Figure 3A we show, as an example, brain renders corresponding to the spatial pattern of the network 9 divided in positive and negative, standing for brain regions that present high correlation and anticorrelation to certain RSNs (DMN and Vis, respectively). Figure 3B shows the level of correlation between each latent mode (N”X”, with “X” varying from 1 to 9) and each RSN indicating with a star the ones that are statistically significant after false discovery rate (FDR) correction. We found that some modes could be associated with the activation of the DMN at the same time with a deactivation of the VIS network (latent networks 6 and 9), while other can be associated with the activation of the Limbic network (N1 and N7). Other latent networks could be described as the activation of Visual areas at the same time of a deactivation of the FP and DMN (N8) Finally, latent networks 4 and 5 could be characterised by the activation of primary/sensory motor function high correlation with VA and DA networks. Interestingly, our association of latent dimensions with activation and deactivation patterns in the brain, shows high similarity with previous results based on non-supervised clustering on brain activity considering the well-known coactivation patterns (CAPs) as is presented by Huang and colleagues (Huang et al., 2020) and with functional gradients analysis (Margulies et al., 2016). Importantly, we replicated this analysis using seven dimensions for the latent space and we found that the networks show similar correspondence with the 7 Yeo RSNs than the results for N=9 (Supplementary Figure 2). We also included in Supplementary material the association between DK62 regions and the Yeo7 functional networks.

**Figure 3.**
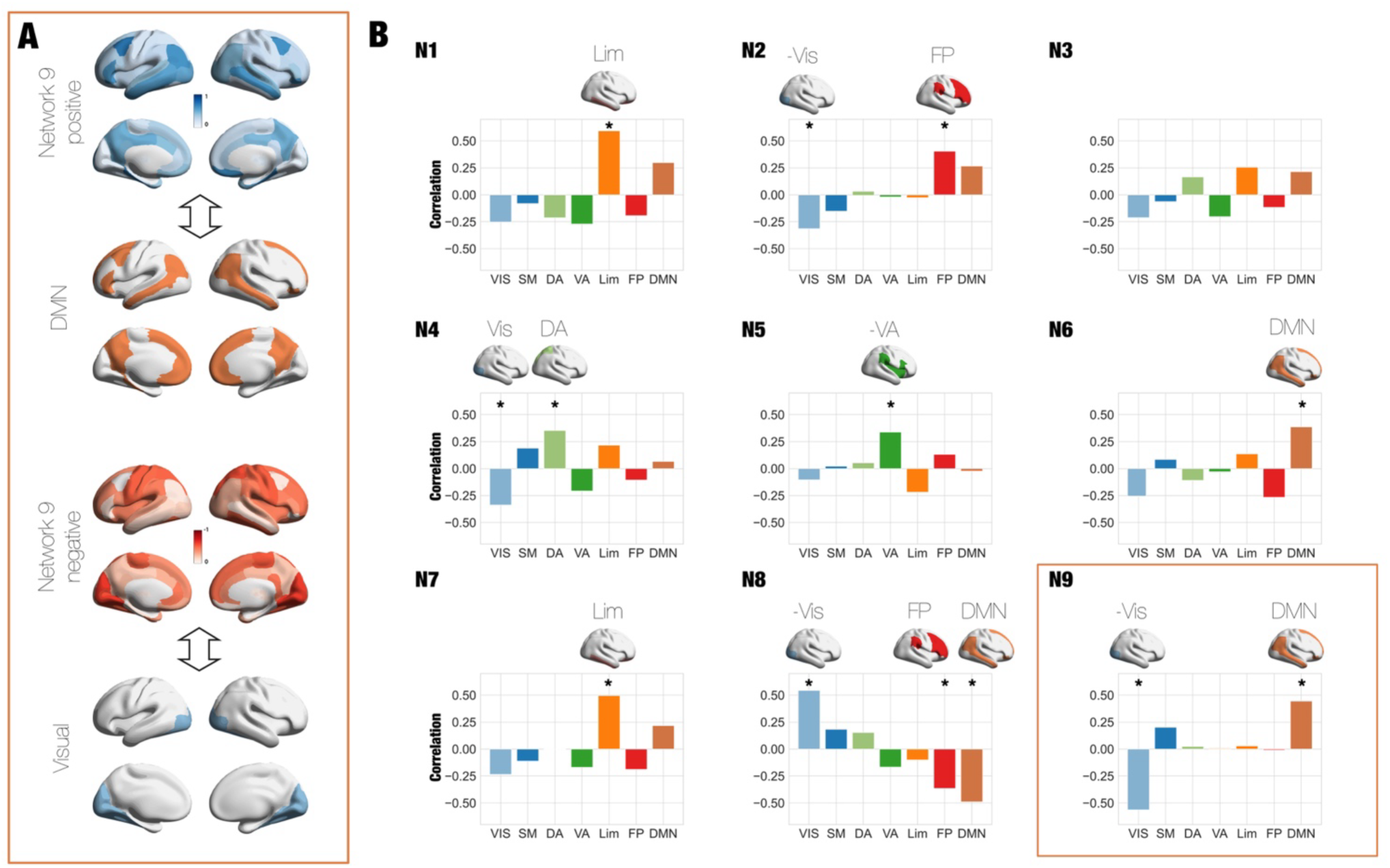
Resting state networks are formed from the latent networks revealed by VAE. **A)** We determined the brain regions associated with each mode. We show as an example the brain renders corresponding to the spatial pattern of the mode 3 divided in mode 3 positive and mode 3 negative, standing for brain regions that present high correlation and anticorrelation to the DMN and Vis network, respectively. **B)** We identified each latent mode with a pattern in the source space of 62 brain regions. We associated each spatial patterns with the Yeo 7 resting state networks by computing the correlation of each pattern with the percentage of belonging of each region to each RSN (* indicates the correlation that are significant after false discovery rate correction). The reference functional brain networks estimated by Yeo and colleagues named: Visual (VIS), Somatomotor (SM), Dorsal Attention (DA), Ventral Attention (VA), Limbic (Lim), Frontoparietal (FP) and Default mode (DMN).

### Reconfiguration of low-dimensional network interaction during cognitive tasks

We then built a model for each task by inferring the effective connectivity between networks in the manifold to obtain the best performance in reproduce 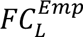 and 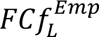 (see Method). Thus, we built 100 models for each task and we assessed each model performance by measuring the correlation between the empirically functional connectivity (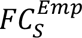) and the latent model-decoded functional connectivity (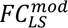) for each task (Figure 3A). For each task, we obtained the LGEC for each one of the 100 models and in Figure 3B we display the average LGEC across model, representing the interaction between networks. Notice that the interactions matrices are capturing the time asymmetry interaction between the networks and consequently are asymmetric. We show how interactions are reconfigured comparing resting-state and social tasks (Figure 3C and D). To quantify how the interaction between low dimensional networks is shaped by cognitive states, we computed the total level of connectivity of each network as the sum of all outcome and income interactions. We found significant differences (Wilcoxon ranksum test, false discovery rate corrected) in the level of TC for almost all networks (except for N7) in the comparison between resting-state and social task (Figure 3C). The comparison between rest and other tasks is displayed in **Supplementary Figure 3** also showing a significant network interaction reconfiguration. We represented with directed graphs the interactions between networks above a threshold (*LGEC_ij_* > 0.1) divided in the outcomes of each network, i.e., how each network impacts to others (**Figure 3D first column**) and the incomes of each network, i.e., how each network is driven by the others, (**Figure 3D second column**). Interestingly, the most important interactions are between networks involving the DMN (N6,8 and 9), which is known to play an important role in the cortical organisation during cognition (Margulies et al., 2016). In particular, the interaction between N2 and N6 is observed during social tasks and not during rest, showing a reconfiguration of that interaction during social.

Finally, we trained a support vector machine (SVM) using the 100 LGEC matrices generated from modelling the manifold for each task presented in Figure 4B. The objective was to determine if the information captured in each matrix could serve as a unique fingerprint for each task. We also trained the SVM classifier using the elements of the empirical functional connectivity (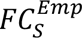), the functional connectivity modelled with the GCAT framework (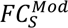) and the LGEC matrices with randomized labels Figure 5A. Our results showed that the best classification accuracy (0.89±0.01) was achieved using the LGEC matrices outperforming the LGEC with randomized classes (0.13±0.03), the 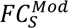 (0.78±0.01) and the 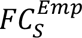 (0.76±0.02) (Figure 5B). In Figure 5C, we also displayed in the rightmost column the confusion matrix obtained for the classification of the 8 labels, the 7 tasks and resting-state.

**Figure 4.**
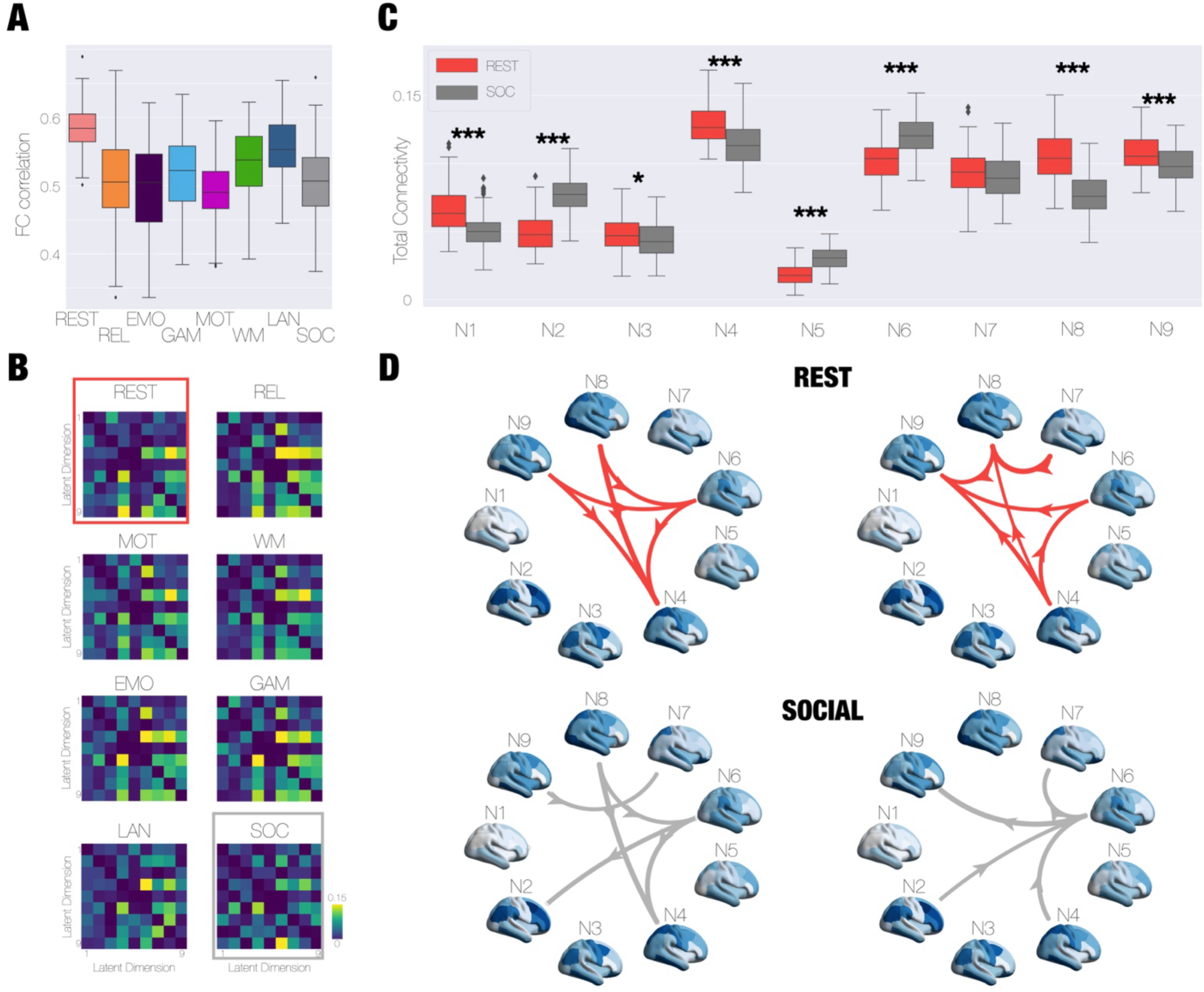
Modelling the low-dimensional manifold network show a flexible reconfiguration of network interaction during cognitive tasks. **A)** We constructed models for each task and evaluated the performance of each model by quantifying the correlation between the 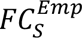 and 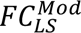 in each case. **B)** The output of each model was the inferred connectivity between the latent variables, called the LGEC, for each task. We show the average LGEC across 100 models for each case. **C)** We computed the total level of connectivity of each network as the sum of all outcome and income interactions. We found significant differences (Wilcoxon ranksum test, false discovery rate corrected) in the level of TC for almost all networks (except for N7) in the comparison between resting-state and social task. (*** means p value<0.001;* means 0.01<p value<0.05) **D)** The highest interactions between networks (above 0.1) represented as a graph. In the left column the outcomes connections, representing how each network drives the others, and in the right column the incomes connection standing for the impact that the rest of the networks have on each network.

**Figure 5.**
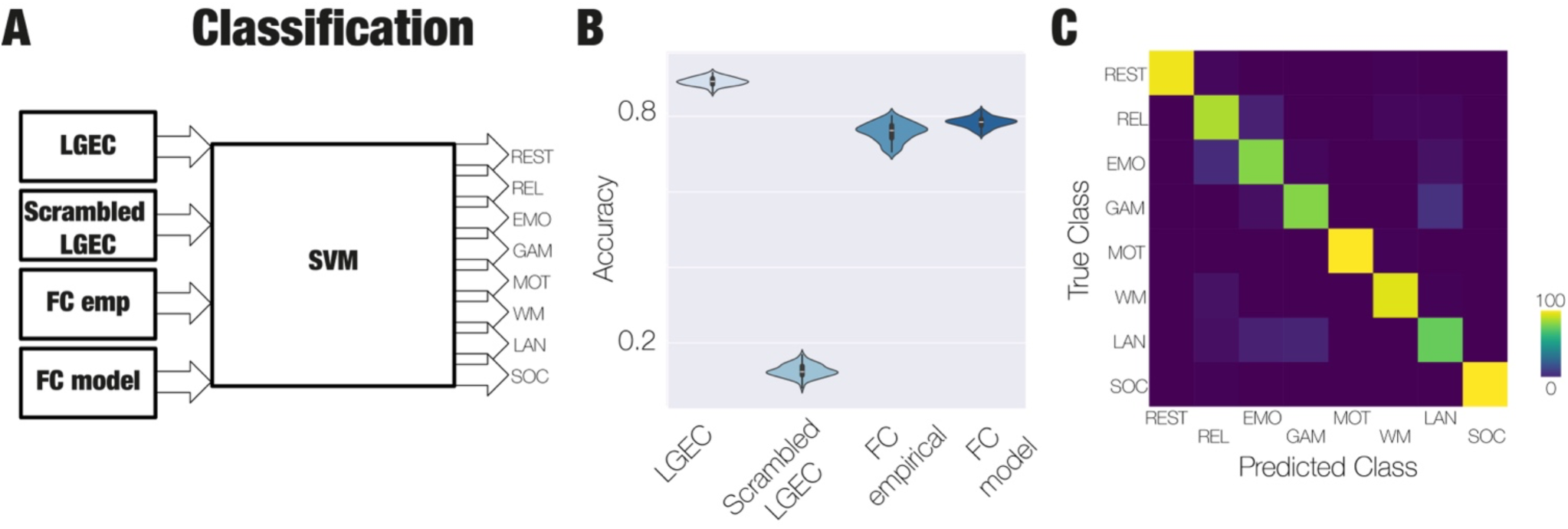
Interaction between low dimensional manifold networks better distinguishes between cognitive tasks. **A)** We trained a support vector machine (SVM) using the 100 LGEC generated by modelling the manifold in each task to evaluate whether the information condensed in each matrix is a fingerprint for each task. We also trained the same SVM classifier but using as an input the elements of the 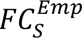, elements of the 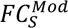 and also the LGEC with scramble labels. **B)** We found that the best classification is reached using the LGCAT (0.89±0.01) compared with the 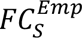 (0.76±0.02), the 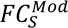 (0.78±0.01) and the scrambled (0.13±0.03). **C)** The confusion matrix obtained for the SVM classifier in the task of distinguish between eight classes, named the resting and the seven cognitive states.

## Discussion

We demonstrated that interactions of low-dimensional manifold networks underlying whole-brain dynamics reveal how large-scale brain organisation is flexible coordinated during cognitive processing. In this sense, our results expand traditional whole-brain models and low-dimensional brain representation analysis (e.g., RSNs (Damoiseaux et al., 2006) or gradients (Margulies et al., 2016)) by modelling the interaction between these low-dimensional networks. To do so, we combined deep learning variational autoencoders and computational modelling to quantitatively infer the interaction between these data-driven obtained networks. Importantly, we showed that generating dynamical models within the subspace in which the low-dimensional manifold is embedded could yield more accurate models in terms of reproducing empirical data compared with modelling the original high-dimensional whole-brain space. Both results are to be expected under the assumption that large-scale dynamics are orchestrated into a reduced number of independent degrees of freedom, which emerge as part of a broader neurobiological principle underlying brain activity. In the following section we discuss the implications of these results, both in terms of their contribution to systems neuroscience and their practical applications to neuroimaging.

### Whole-brain fMRI dynamics are intrinsically low-dimensional

Several past studies investigated the compressibility of resting state fMRI data using linear methods such as principal (Carbonell et al., 2011; Margulies et al., 2016) and/or independent component analysis (Damoiseaux et al., 2006), revealing a set of canonical RSN overlapping with distinct neuroanatomical regions subserving specific cognitive functions (Smith et al., 2009). Other methods, such as the clustering of co-activation patterns (CAPs) (Liu & Duyn, 2013) results in similar information. These networks also show alterations in certain neuropsychiatric disorders (Greicius, 2008), and contain information that can be used to predict and decode the contents of ongoing consciousness and cognition (Cole et al., 2016); moreover, the neurobiological underpinnings of the RSN are highlighted by multimodal imaging studies revealing that their activity correlates with different frequency bands measured using simultaneous EEG (Mantini et al., 2007). As shown **in** Figure 3, the networks representing cognitive processing revealed using autoencoders are related to the RSN, although this relationship is not a simple one-to-one mapping. Instead, these networks are a combination of different RSN, indicating that whole-brain dynamics can be spanned by different sets of functional networks. We also observed the same behaviour when considering seven latent dimensions, showing despite the number of networks is the same there not a one-to-one mapping between both set of networks. Despite the mapping between RSN and networks, the latter have distinct properties due to the nature of the generative autoencoder algorithm. Besides its capacity to capture nonlinear relationships between the networks in the manifold and the high-dimensional data, autoencoders are generative, which is of key importance to translate the dynamics in the latent space towards that of resting-state high-dimensional fMRI data. Crucially, this generative capacity allows us to create dynamical models of interactive networks fitted to the whole-brain dynamics revealing that changes in these interactions are reorganised during cognitive processing.

### Large-scale manifolds and cognition

The use of dimensionality reduction techniques is a common strategy in statistics and machine learning, applied with the purpose of simplifying the description of the data by performing a reasonable grouping of the variables (e.g. clustering) or expressing them as a linear or nonlinear combination of a reduced set of components (e.g. ICA, PCA) (Pang et al., 2016). The application of these methods to neural activity recordings is not only useful in terms of data processing, as expected, but also reveals important features of how the brain represents information relevant to cognition and behaviour (Chung & Abbott, 2021; Vidaurre et al., 2018). Research conducted using a variety of methods and experimental organisms shows converging evidence of the manifold organization of neural activity, where the manifold dimensions can be parametrized by variables with correspondence to behavioural observations. Examples include the representation of movement parameters in the motor cortex (Gallego et al., 2017), the mapping of spatio-temporal coordinates to the hippocampal cortex during navigation (Derdikman & Moser, 2010), and the encoding of sensory (e.g. visual) (Stringer et al., 2019), olfactory (Chandak & Raman, 2023)) information. While these manifolds can change in response to learning (Nieh et al., 2021), they are generally stable (Gallego et al., 2020), and it is believed that they relate to the network of synaptic connections and their strengths; however, the relationship between manifolds and structural features remains difficult to demonstrate (Langdon et al., 2023). Our results suggest that large-scale brain activity also benefits from the organization into low-dimensional manifolds. Indeed, the corresponding dimensions (i.e. networks) may parametrize the engagement of different cognitive functions in spontaneous mentation. Thus, the organization of brain activity in the form of manifolds appears to occur across scales, where coarser scales index the participation of large assemblies of cells, whose dynamics may also be understood in terms of a low-dimensional space.

### Interaction of networks in the manifold and cognition

As we discussed above, low-dimensional representation of brain activity has been widely used to understand large-scale brain organisation during cognition. Here, we focused on modeling the low-dimensional space that allows us not only to enhance the fitting quality, eliminating the need to make the model replicate redundant or noisy data from the high-dimensional space, but also allow us to infer the interaction between the networks. Consequently, we went beyond the pure low-dimensional representation and focused how the interactions of these networks embedded in that low-dimensional manifold are reconfigured with task activity. We found that these interactions are reorganised during cognition and the classification of tasks from the HCP is optimal when based on the low-dimensional network representation, suggesting that these manifold dimensions are not arbitrary but encode organizational principles of brain function of cognitive processing instead (Figure 5). We also found the most important interactions are between networks involving the DMN (N6,8 and 9), which is known to play an important role in the cortical organisation during cognition and primary/sensory motor function such as the latent network 4 (Margulies et al., 2016) (Figure 4). Overall, our results not only broadcast the relevance of the low-dimensional brain representation but also the interaction between networks in that space provides crucial information for understanding cognitive processing.

### Possible disruptions in disease and unconsciousness

As in the case of RSN, we expect that networks and its interactions are sensitive to different neurological and psychiatric conditions. If these networks reflect the representation of ongoing cognition and consciousness in large-scale patterns of brain activity, we could expect the most salient alterations in disorders leading to severe impairments in these domains. An example is the case of brain injured patients who show a state of stable and persisting unresponsiveness, which can be interpreted as a lack of conscious thought (Schiff et al., 2014). Diminished or absent conscious content can also be present in certain transient states, either spontaneously occurring (such as deep sleep) or induced by pharmacological means (Brown et al., 2010). Previous research shows that a low-dimensional representation of brain dynamics captures the differences between the aforementioned states and conscious wakefulness, the former showing a reduced repertoire of states visited during the recording, and less structured and complex transition between (Rué-Queralt et al., 2021) these states (Varley et al., 2021). Future studies should adopt the framework developed here to investigate different pathological and physiological brain states, with the purpose of obtaining complementary information on the landmark features of brain dynamics during health and disease.

### Modelling the coupled modes dynamics

Our work provides novel methodological perspectives for the analysis of large-scale neuroimaging data. The majority of whole-brain modelling studies begins by adopting a certain parcellation of the human brain into regions of interest (ROI). Assuming the anatomical connectivity between ROI is known, it is then possible to couple the local equations and simulate whole-brain dynamics. Our work suggests that this methodology may be suboptimal since brain dynamics are better represented by modes which span multiple ROIs; instead, models should strive to capture the relationship between networks. While the anatomical connectivity between the distributed sets of regions spanning the neural nodes is ill-defined, our result shows that equivalent results can be obtained using the effective connectivity. The generative capacity of autoencoders represents another attractive characteristic of our work, as it allows us to create new data following similar statistics as the empirical recordings, with potential applications to machine learning and data augmentation. Importantly, our approach can be applied to different neuroimaging modalities such EEG/MEG allowing to create models of the low-dimensional networks.

### Limitations and future work

Our work presents some limitations that should be addressed in future studies. Perhaps the most salient one stems from our use of fMRI data, with known limitations to its biological interpretability. This should be assisted by other sources of information capable of complementing the recordings and shedding light on the origin of the neural models. Also, the use of improved MRI machines with higher fields could transcend the spatial organization of the cortex into maps, allowing researchers to focus on layer-specific contributions. Finally, the set of tasks to be distinguished was relatively small. A larger repertoire of conditions should be included to tackle questions concerning the specificity of the underlying modes to classify pathological and physiological brain states.

### Conclusions

We demonstrated that interactions of low-dimensional manifold networks underlying whole-brain dynamics reveal how large-scale brain organisation is flexible coordinated during cognitive processing. Importantly, our results not only change the focus from brain regions to low-dimensional networks but also from network description towards network interaction. We quantified the interactions between these networks combining deep learning autoencoders and dynamical modelling, leading to better results than the ones obtained in the original high dimensionality space. We conclude that standard parcellations of brain activity are prone to overlook the underlying manifold organization of fMRI, and that future studies should attempt to characterize and model brain states adopting the perspective of interactive networks.

## Acknowledgments

Y.S.P is supported by NEMESIS project (ref. 101071900) funded by the EU ERC Synergy Horizon Europe. M.L.K. is supported by the Center for Music in the Brain, funded by the Danish National Research Foundation (DNRF117), and Centre for Eudaimonia and Human Flourishing at Linacre College funded by the Pettit and Carlsberg Foundations.

ET is supported by grants FONCYT-PICT (2019-02294), CONICET-PIP (11220210100800CO), and ANID/FONDECYT Regular (1220995). G.D. was supported by a number of sources: NEMESIS project (ref. 101071900) funded by the EU ERC Synergy Horizon Europe; AGAUR research support grant (ref. 2021 SGR 00917) funded by the Department of Research and Universities of the Generalitat of Catalunya; project PID2022-136216NB-I00 financed by the MCIN /AEI /10.13039/501100011033 / FEDER, UE., the Ministry of Science and Innovation, the State Research Agency and the European Regional Development Fund.

## Author contributions

Y.S.P, M.L.K, E.T. and GD designed the research. Y.S.P, E.T. and GD conducted the research. Y.S.P., M.L.K.,G.D and E.T. analyzed and interpreted the results. M.L.K curate the data. Y.S.P, M.L.K., E.T. and GD wrote the manuscript and made figures. Y.S.P. analyzed the data. E.P.B & Y.S.P, created and published the code. G.D and E.T, supervised the research. All authors provided analytic support. All authors edited the manuscripts.

## Methods

### Neuroimaging Ethics

The Washington University–University of Minnesota (WU-Minn HCP) Consortium obtained full informed consent from all participants, and research procedures and ethical guidelines were followed in accordance with Washington University institutional review board approval.

### Neuroimaging Participants

The neuroimaging dataset used in this investigation was obtained from the HCP (Human Connectome Project) March 2017 public data release. From the total of 1,003 participants available in the release, a sample of 1000 to the first part when only considered rest and 990 participants was chosen for the second part due to not all participants performing all tasks.

The HCP task battery comprises seven tasks, namely working memory, motor, gambling, language, social, emotional, and relational tasks. Detailed descriptions of these tasks can be found on the HCP website (Barch et al., 2013). The tasks were intentionally designed by HCP to cover seven major cognitive domains, which aim to capture the diversity of neural systems: 1) Visual, motion, somatosensory, and motor systems; 2) Working memory, decision-making, and cognitive control systems; 3) Category-specific representations;4) Language processing; 5) Relational processing;6) Social cognition; 7) Emotion processing.

Apart from the resting-state scans, all 1,003 participants completed all tasks in two separate sessions. The first session included working memory, gambling, and motor tasks, while the second session involved language processing, social cognition, relational processing, and emotion processing tasks.

### Neuroimaging extraction of functional time series

The processing and extracting data from both resting-state and task-based fMRI datasets is detailed in the HCP (Human Connectome Project). The preprocessing steps, described in detail on the HCP website, utilized standardized methods from FSL (FMRIB Software Library), FreeSurfer, and the Connectome Workbench software. The process included correcting spatial and gradient distortions, addressing head motion, intensity normalization, bias field removal, registration to the T1-weighted structural image, and transformation to the 2-mm Montreal Neurological Institute (MNI) space. The FIX artefact removal procedure (Schröder et al., 2015; Smith et al., 2013) and ICA+FIX processing (Arslan et al., 2018; Griffanti et al., 2017) were employed to remove structured artefacts and head motion parameters, respectively.

The preprocessed time series data of all grayordinates were in HCP CIFTI grayordinates standard space and available in surface-based CIFTI files for each participant during resting-state and each of the seven tasks. A custom-made MATLAB script using the ft_read_cifti function from the FieldTrip toolbox was used to extract the average time series for all the grayordinates in each region of the each parcellation used, as defined in the HCP CIFTI grayordinates standard space. The BOLD timeseries were filtered using a second-order Butterworth filter with a frequency range of 0.008–0.08Hz and extracted the first and last 5% volumes to avoid the borders effect of the filtering.

### Parcellations

The neuroimaging data underwent processing using the three standard parcellations with different amount of brain cortical regions: i) the Mindboggle-modified Desikan-Killiany parcellation, called DK62 (Desikan et al., 2006); ii) the Schaefer considering 500 regions; and iii) the Schaefer parcellation considering 1000 regions This DK62 parcellation consists of a total of 62 cortical regions, with 31 regions in each hemisphere, as defined by Klein and Tourville in 2012. Consequently, the DK80 parcellation contains a total of 62 regions, precisely defined in the common HCP CIFTI grayordinates standard space. Then, the Schaefer 500 and 1000 parcellation was also used with 250 and 500 cortical regions per hemisphere respectively, and well defined in the HCP CIFTI. In this parcellation(Schaefer et al., 2018)

### Variational Autoencoders architecture and training

The VAE consists of the same three structures as the AE, but it incorporates certain differences that give VAEs generative properties. To guarantee this feature, during training, errors were propagated through gradient descent to minimize a loss function consisting of two terms. The first term is a standard reconstruction error, computed from the units in the output layer of the decoder. The second term is a regularization term, calculated as the Kullback-Leibler divergence between the distribution in the latent space and a standard Gaussian distribution. This regularization term ensures continuity and completeness in the latent space, meaning that similar values are decoded into similar outputs and that these outputs represent meaningful combinations of the encoded inputs. Our encoder network is composed of a deep neural network with rectified linear units as activation functions and two dense layers. This network funnels down to a low-dimensional variational layer, spanning the latent space explored from dimensions 5 to 12. The encoder network applies a nonlinear transformation to map the values of each time pattern to Gaussian probability distributions within the latent space. Conversely, the decoder network replicates the architecture of the encoder, generating reconstructed temporal patterns by sampling from these distributions. We trained the VAE with time-by-time spatial patterns for the training set creating from the 90%/10% (training/testing) resting state HCP participants full dataset. The network training procedure consisted of batches with 128 samples and 50 training epochs using an Adam optimizer and the training consists of error backpropagation via gradient descent to minimize a loss function composed the two terms.

In the last section of this work, we extended the framework by including to resting state (REST) data the recordings from 7 cognitive tasks from HCP (Social [SOC], Language [LAN], Working Memory [WM], Motor [MOT], Gambling [GAM], Emotion [EMO], Relational [REL]) to find the low-dimensional manifold that represent the dynamics which supports cognitive processing. We repeated the procedure as explained above, but in this case, we train the VAE using the time-by-time spatial patterns from the full data by concatenating the rest condition and the 7 tasks. We fixed the latent dimensions to 9 and we used the DK62 parcellation based on our previous results to build models in the latent space.

### Modes coupled model and LGEC inference

We modelled the local dynamics of each latent neural mode as a Stuart-Landau oscillator (a dynamical system that can be expressed as the normal form of a supercritical Hopf bifurcation), this model presents the capability to shift from noisy to oscillatory dynamics. Importantly, this kind of applied to the whole-brain dynamics have been able to replicate key aspects of brain dynamics observed in electrophysiology, magnetoencephalography(Cabral et al., 2014) and fMRI(Deco et al., 2017; Ipiña et al., 2020; Yonatan S Perl et al., 2022). Specifically, given a parcellation that number *M* of modes, the whole-brain dynamics of the coupled modes can be expressed by the local dynamics of *M* Stuart-Landau oscillators connected via the connectivity matrix ***C***, which is defined by:

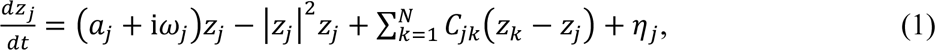

where the complex variable *Z_j_* denotes the state (*Z_j_* = *x_j_* + i*y_j_*) of mode *j*, *η_j_* is additive uncorrelated Gaussian noise with variance *σ*^2^ (for all *j*), *ω_j_* is the intrinsic mode frequency. The intrinsic frequencies *ω_j_* (which lie in the 0.008–0.08Hz band) were estimated from the empirical data as the averaged peak frequencies of the narrowband the signals of the different modes. Finally, the mode’s bifurcation parameter is *a_j_*which defines the dynamics of the mode with respect to the Hopf bifurcation. This bifurcation is characterised by bifurcation parameter (*a*) that induces qualitative changes in the dynamical behaviour when the system crosses the bifurcation point (*a* = 0). In the supercritical regime (*a* > 0,), the oscillator generates self-sustained oscillations, whereas in the subcritical regime (*a* < 0), the dynamics converge to a fixed point. Importantly, the addition of noise to the model causes the dynamics near the bifurcation point (*a*∼0) to stochastically transition between both regimes, resulting in oscillations with complex amplitude modulations. Previous studies have demonstrated that this regime represents the optimal point for fitting empirical data in whole-brain models (Deco et al., 2017; Ipiña et al., 2020; Yonatan S Perl et al., 2022). We fix the bifurcation parameter of each mode in that regime (*a* = −0.02).

We fit ***C*** such that the model optimally reproduces the empirically correlation matrix in the latent space 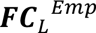 (i.e., the pairwise correlation between the latent modes) and the empirical forward time-shifted correlation between the latent modes 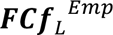(*τ*). We selected the parameter *τ* = 3 which led to a decrease in the averaged autocorrelation also based on previous works(Deco et al., 2022). We note that fitting the time-shifted correlations can lead to asymmetries in the connectivity ***C***, which, in turn, can produce non-equilibrium dynamics. Using a heuristic pseudo-gradient algorithm, we proceeded to update the ***C*** until the fit is fully optimised. More specifically, the updating uses the following form:

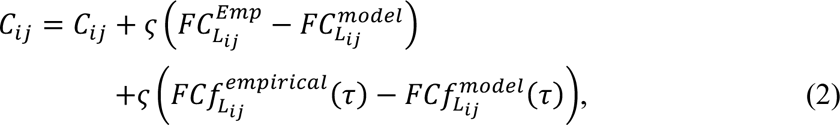

The model was run repeatedly with the updated ***C*** until the fit converges towards a stable value. We initialized ***C*** as a zero matrix of dimension *MxM* (the number of latent modes) and we update all the connections from this matrix (in either hemisphere). For the Stuart-Landau model, we used 𝜍 = 0.00001 and continue until the algorithm converges. We generate 100 model by repeating this procedure to fit the average empirical matrices, which was obtained by averaging set of 10 (9) participants for the resting state (resting state plus tasks). In this way we manage to improve the fitting of the models. In each model, for each iteration we average over 10 (9) simulations as we made for the empirical data. Overall, we use the term Latent Generative Effective Connectivity (LGEC) for the optimised ***C***, which reflects the connectivity between the modes.

### Obtaining sources networks by decoding the latent modes

We created a set of surrogate signals by introducing standard Gaussian noise into one latent dimension while keeping the other eight dimensions devoid of any signal. We repeated this process, varying the dimension with noise and the dimensions without any signal, and we called N “X” with X from 1 to 9 to each one. Subsequently, we decoded each of these nine sets of surrogate latent signals, mapping them from the latent dimensions to the 62 brain regions of the original empirical data for each case. Next, we examined the regions in the source space that were activated in response to the presence of noise in each latent dimension. We quantified the variance of the decoded time signals to evaluate the statistical response of the activity associated with each latent dimension. This analysis allowed us to obtain one spatial pattern in the empirical original space associated to each latent dimension.

### Support vector machine classifier

We use a SVM with Gaussian kernels as implemented in the MATLAB function fitcecoc. The function returns a full, trained, multiclass, error-correcting output codes (ECOC) model. This is achieved using the predictors in the input with class labels. The function uses K(K − 1)/2 binary SVM models using the one-versus-one coding design, where we used K = 8 as the number of unique class labels. In other words, the SVM had as inputs the elements of the LGEC obtained in each of the 100 models for each one of the 7 tasks and the resting-state condition. The output was eight classes corresponding to the conditions (rest and seven tasks). We subdivided into 90% training and 10% validation, repeated and shuffled 100 times. For the classification using the functional connectivity in the original source space, we average the FC across 9 participants, as we did with the LGEC computation, and use as an input the upper diagonal elements of the matrices.

### Statistical analyses

We applied the Wilcoxon rank-sum method to test the significance and we applied the False Discovery Rate (FDR) at the 0.05 level of significance to correct multiple comparisons(Hochberg & Benjamini, 1990).

### Data availability

The dataset utilized in this study originated from an independent publicly accessible collection of fMRI data. In this case, we opted for a sample comprising 1003 participants, drawn from the Human Connectome Project’s March 2017 public data release (https://www.humanconnectome.org/study/hcpyoung-adult)

### Code availability

The code used in this paper is deposited in https://github.com/yonisanzperl/ModellingManifoldModes. The software dependencies are MATLAB (2018b); Python (3.6) and Keras. From time to time, the code might be updated.

